# Comparative genomic and crystal structure analyses identify a collagen glucosyltransferase from *Acanthamoeba Polyphaga Mimivirus*

**DOI:** 10.1101/2022.05.07.491030

**Authors:** Wenhui Wu, Jeong Seon Kim, Stephen J. Richards, Christoph Buhlheller, Aaron O. Bailey, William Russell, Tiantian Chen, Tingfei Chen, Zhenhang Chen, Bo Liang, Mitsuo Yamauchi, Houfu Guo

## Abstract

Humans and *Acanthamoeba Polyphaga Mimivirus* share numerous homologous genes, including collagens and collagen-modifying enzymes. To explore the homology, we performed a genome-wide comparison between human and mimivirus using DELTA-BLAST (Domain Enhanced Lookup Time Accelerated BLAST) and identified 52 new mimiviral proteins that are homologous with human proteins. To gain functional insights into mimiviral proteins, their human protein homologs were organized into Gene Ontology (GO) and REACTOME pathways to build a functional network. Collagen and collagen-modifying enzymes form the largest subnetwork with most nodes. Further analysis of this subnetwork identified a putative collagen glycosyltransferase R699. Protein expression test suggested that R699 is highly expressed in *E coli*, unlike the human collagen-modifying enzymes. Enzymatic activity assays showed that R699 catalyzes the conversion of unique galactosylhydroxylysine within the GXXXUG motif (U=galactosylhydroxylysine) to glucosylgalactosylhydroxylysine on collagen using uridine diphosphate glucose (UDP-Glc) as a sugar donor, suggesting R699 is a mimiviral collagen galactosylhydroxylysyl glucosyltransferase (GGT) with defined substrate specificity. Structural study of R699 produced the first crystal structure of a collagen GGT with a visible UDP-Glc. Sugar moiety of the UDP-Glc resides in a previously unrecognized pocket. Mn^2+^ coordination and nucleoside-diphosphate binding site are conserved among GGT family members and critical for R699’s collagen GGT activity. To facilitate further analysis of human and mimiviral homologous proteins, we presented an interactive and searchable genome-wide comparison website for quickly browsing human and *Acanthamoeba Polyphaga Mimivirus* homologs, which is available at RRID Resource ID: SCR_022140 or https://guolab.shinyapps.io/app-mimivirus-publication/.

## Introduction

In humans, collagens represent the most abundant protein family forming the extracellular matrix to support and regulate cells and to maintain tissue form and stability (1,2). At least 28 members of collagen family have been identified and each member likely carries out specific functions (3). Fibrillar type I collagen is the most abundant member in the family providing most connective tissues with mechanical strength. Type I collagen is a heterotrimeric molecule composed of two a1 and one a2 chains, and it consists of three domains: N-and C-terminal non-triple helical domain (N- and C-telopeptides) and the central triple helical domain. As the major collagen component of the basement membrane, type IV collagen is a heterotrimeric network-forming collagen underlying epithelial and endothelial cells and functioning as a barrier between tissue compartments. Type IV collagen contains the N-terminal 7S, a central triple-helical domain, and the globular C-terminal NC1. To perform their functions, collagens acquire a series of specific post-translational modifications (PTMs) during biosynthesis (4). Collagen prolyl 4-hydroxylation catalyzed by collagen prolyl 4-hydroxylases is critical for stabilizing the triple-helical structure of collagens (4). Prolyl 3-hydroxylases produce prolyl 3-hydroxylation and defects in this modification are associated with recessive osteogenesis imperfecta (5). A series of lysine (Lys) PTMs of collagens are critical for the stability of collagen fibrils. In the cells, Lys residues in the sequences of X-Lys-Gly (helical domain) and X-Lys-Ala/Ser (telopeptides) can be hydroxylated by lysyl hydroxylases 1-3 (LH1-3) to form 5-hydroxylysine (Hyl) (6). It is generally accepted that LH1 is the main LH for the helical domain and LH2 for the telopeptides. Certain Hyl residues in the collagen helical domain are galactosylated by glycosyltransferase 25 domain containing 1 and 2 (GLT25D1 and GLT25D2) and then glucosylated by lysyl hydroxylases to form a unique Hyl-*O*-linked glycosylation with a mono-or di-saccharide (7-12). LH3 is the major galactosylhydroxylysyl glucosyltransferase (GGT) catalyzing collagen glucosylation (8), although it has been shown to be a multifunctional enzyme (13). Collagen prolyl and lysyl PTMs are tightly regulated during the development and their alterations lead to various diseases (4,14). For instance, mutations in the gene encoding LH2 result in Bruck syndrome II a rare osteogenesis imperfect with joint contracture, but hyper LH2 activity contributes to fibrosis and cancer growth and metastasis (15-23).

Besides multicellular animals, collagen-like proteins and collagen-modifying enzymes are highly conserved across species and have been found in certain fungi, bacteria, and viruses such as mimivirus (24-27). Since the initial release of the mimiviral genome (28), studies have identified 7 mimiviral collagen genes and 2 mimiviral collagen-modifying enzyme genes that encode three enzymes, including collagen prolyl hydroxylase, collagen lysyl hydroxylase, and collagen hydroxylysyl glucosyltransferase (26,29). Structural and functional studies of mimiviral collagen lysyl hydroxylase provide insights into functions of the human collagen-modifying enzymes (12,30). Since collagen is widely used for tissue and biomaterial engineering, efforts have been made to express recombinant collagens using different expression systems (31-33). Interestingly, a hydroxylated human collagen III fragment has been produced in *E coli* by coexpressing it with mimiviral collagen prolyl and lysyl hydroxylases (29). However, glycosylated human collagen is still unable to be produced in the bacterial expression system, at least due in part to the difficulty of expressing active human collagen glycosyltransferases in bacteria.

Mimivirus is the first giant virus discovered and is the best-characterized virus in the family (28). The initial mimiviral genome sequencing effort identified 917 protein-encoding genes (28). These genes play diverse functions in nucleotide and protein biosynthesis, including DNA replication, repair, transcription and translation (34-38). This effort also identified enzymes involved in various PTMs including 11 glycosyltransferases (28). As the sequencing technique advances, a later sequencing analysis identified 75 new genes and pushed the mimiviral genes to exceed 1000 (39). More recent work identified citric acid cycle and β-oxidation pathway genes in the mimiviridae family (40,41). Since the release of the mimiviral genome sequence and the first search for its homology to other species (28), more than 50 mimiviral proteins have been expressed and characterized (Supplemental Table. 1), which provides valuable insights into virology and raises questions regarding the definition of viruses.

To facilitate the further study of mimiviral homologous proteins, a systematic search of mimiviral homologous proteins in humans was performed. We compared human and mimiviral proteins at the genome-wide level using the DELTA-BLAST (Domain Enhanced Lookup Time Accelerated BLAST) (42). Besides the initially identified 194 mimiviral ORFs that shared homology with known proteins mainly involving in DNA and protein metabolism, we found 52 new mimiviral ORFs that show similarity to human proteins. Eight mimiviral collagen-like proteins (L71, L668, L669, R196, R238, R239, R240, and R241) and 4 putative mimiviral collagen-modifying enzymes (L230, L593, R655, and R699) were identified. To validate the results, we expressed a putative mimiviral collagen glycosyltransferase R699, characterized its activity, and solved its crystal structure. Moreover, we established an interactive and searchable genome-wide comparison tool. This user-friendly website helps users quickly browse the protein sequence homology between humans and mimivirus at the genome-wide level for querying new homologs and generating new hypotheses.

## Results and Discussion

### Human and mimivirus homology

We used 979 mimiviral ORFs to query human homologs in human non-redundant protein sequences using DELTA-BLAST. When using e-value <= 0.01, hit span >= 35 aa, % identical sequences >= 0.25 as cutoffs, 322 queries resulted in at least 1 hit with 41521 hits in total. The search found 4123 unique human RefSeq records in total. Further analysis showed that the 4123 unique human RefSeq records are from 1236 unique human proteins (Supplemental Table. 2). To identify mimiviral protein homologs using human protein queries, we used 4123 unique human RefSeq records that we identified in the first round of analysis to search for mimiviral homologs. Using the same filtering standard, we found that 3325 RefSeq queries or 1049 human proteins result in at least 1 hit and 58011 hits in total (Supplemental Table. 3). This search found 307 mimiviral protein sequences, 265 of which overlap with the 322 mimiviral queries that we started with. Of these 265 mimiviral queries, 52 of them are newly identified (Supplemental Table. 4). The most common motif shared by humans and mimivirus is ankyrin repeats. Besides the 48 mimiviral ankyrin repeats previously identified, our analyses added 33 more mimiviral proteins with ankyrin repeats.

To identify the enriched pathways conserved between humans and mimivirus, we performed GO and REACTOME pathway analysis using an adjusted p-value <= 0.05 as a cutoff (Supplemental Table. 5-8). GO pathways were organized based on cellular component, molecular function, and biological process (Supplemental Table. 5-7). Functional enriched GO and REACTOME pathways were then used to build gene ontology networks and visualized using *Cytoscape3.8.2* (**Fig. 1A**). This analysis identifies 52 clusters involved in endocytosis, ubiquitination, DNA and collagen metabolism. The largest cluster with the most nodes is the network forming collagen composed of collagen and collagen-modifying enzymes (**Fig. 1A**). Collagen-related pathways rank high in both GO and REACTOME pathway analyses (**Fig. 1B** and Supplemental Table. 5-8).

**Figure 1.**
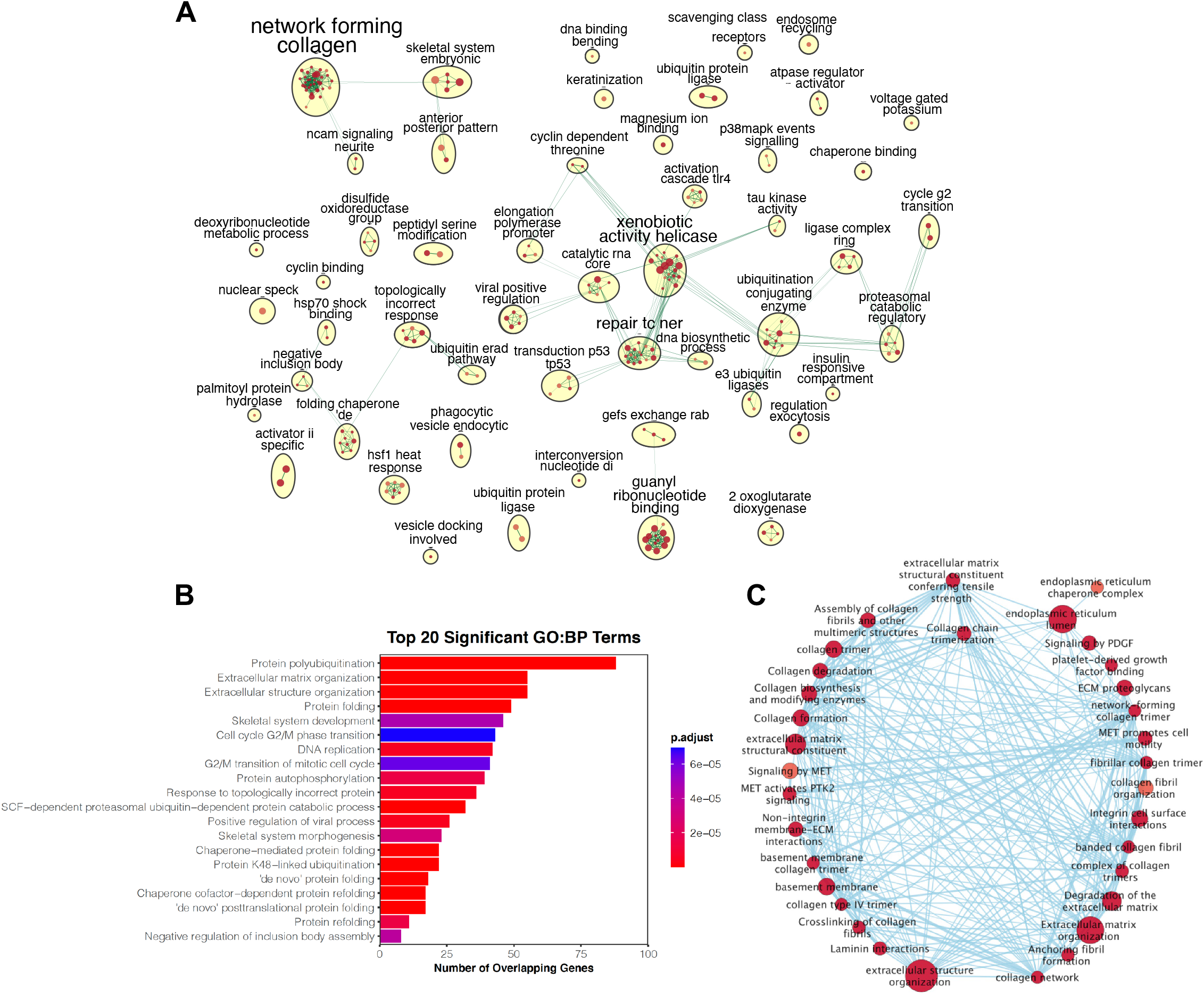
Comparative genomic analysis of human and mimivirus. A, Functional enriched Gene Ontology (GO) and REACTOME pathways that are shared by human and mimivirus. We performed a genome-wide search of homologous genes in human genome using mimiviral proteins as queries. This search identified 4123 unique human RefSeq records that were organized into GO and REACTOME pathways. Gene ontology networks were built based on the results from pathway analysis and visualized using *Cytoscape3.8.2*. Collagen related pathways form the largest subnetwork with the most nodes. B, Top 20 GO enriched biological processes were shown. Collagen-related pathways are the major components of Extracellular matrix organization and Extracellular structure organization. C, Subnetwork analysis shows the organization of collagen-related pathways.

### Homology in collagen and collagen-modifying enzymes

We performed subnetwork analysis on network forming collagen since it is the largest subnetwork with most nodes (**Fig. 1C**). Subnetwork analysis showed that the human protein hits we identified using mimiviral queries are involved in collagen biosynthesis and assembly. Our search identified 8 mimiviral collagens (L71, L668, L669, R196, R238, R239, R240, and R241) and 4 mimiviral collagen-modifying enzymes (L230, L593, R655, and R699). Of these 12 collagen and collagen-modifying enzyme genes, the identity of R238, R655, and R699 has not been revealed. Sequence homology analyses suggest that R238 is a collagen-like protein while R655 and R699 are putative collagen glycosyltransferases. Eight of the mimiviral collagen-related proteins identified by the initial search were not correctly annotated. For instance, 4 mimiviral collagen-like proteins (R196, L669, R239, and R241) had been annotated as PPE-repeat proteins and 2 putative collagen-modifying enzymes (L230 and R655) as LPS biosynthesis enzymes (28). Of these mis-annotated mimiviral proteins, L230 has been expressed and characterized as collagen lysyl hydroxylase and collagen hydroxylysyl glucosyltransferase (26). Structural and mutagenesis analyses suggest that L230 lysyl hydroxylase domain forms a Fe^2+^-stabilized tail-to-tail homodimer (30), similar to human lysyl hydroxylase 2. For the mimiviral proteins that were not annotated during the initial release of the mimiviral genome, L71 has been confirmed to be a type of mimiviral collagen that may play a role in the pathogenesis of arthritis in humans (43). L593 was shown to hydroxylate human type III collagen proline residue and was used to generate a human recombinant collagen III fragment in a bacterial expression system (29). These results validate our search of the viral collagen and collagen-modifying enzymes, suggesting that our analyses are robust and relevant. No lysyl oxidase or transglutaminase was identified, suggesting that mimiviral collagens are either crosslinked by host enzymes or not crosslinked at all.

### R699 has collagen GGT activity

Of the two unstudied putative collagen-modifying enzymes, R655 has been linked to be a putative mimiviral glycosyltransferase (26) while little is known about R699. As a result, we picked R699 for further biochemical and structural analyses. We hypothesized that R699 is a collagen GGT because R699 shows a higher sequence similarity to human collagen GGTs than collagen hydroxylysyl galactosyltransferases (**Fig. 2A**). To test whether R699 encodes a protein, we synthesized the R699 gene and expressed it in *E coli*. We found that R699 produces a stable protein with a yield of ∼10 mg per liter of *E coli* culture (**Fig. 2B**). Highly purified R699 protein was obtained after a 4-step purification procedure (**Fig. 2B**). Given the moderate sequence similarity (30% amino acid sequence identity) between R669 and human GGTs, we speculated that R699 might function as a collagen GGT. To test this possibility, we reacted R699 with galactosylhydroxylysine (**Fig. 2C-E**) or deglucosylated type IV collagen substrate (**Fig. 2F-I**) using UDP-Glc and 3 other sugar donors. Deglucosylation of type IV collagen was generated as previously described (**Fig. 2F** and supplemental Fig. 1) (12). Under these conditions, R699 showed robust activity with UDP-Glc but no other sugar donors (**Fig. 2C and 2G**). We also performed the GGT enzymatic activity assay using UDP-[UL-^13^C_6_]-Glc as a sugar donor to confirm the glucosylation events and map the molecular loci of glucosylation. The modification of galactosylhydroxylysine was quantified and confirmed by liquid chromatography-mass spectrometry (LC-MS) analysis (**Fig. 2D and 2E**). The molecular loci of glucosylation on type IV collagen were mapped with LC-MS (**Fig. 2H and 2I** and Supplemental Table. 9-11). Unlike LH3 that promiscuously glucosylates galactosylhydroxylysine residues within a variety of sequence contexts, R699 preferentially modifies a unique substrate peptide sequence with the non-canonical collagen motif GXXXUG (U=galactosylhydroxylysine), which results in the major differences in consensus motif recognized by R699 and LH3 (**Fig 2J**). These results suggest that R699 is a collagen GGT with a unique substrate specificity. Given that mimiviral L230 functions as a collagen LH and a hydroxylysyl glucosyltransferase to produce peptidyl glucosylhydroxylysine (26), it is tempting to test the possibility of R699 modifying peptidyl glucosylhydroxylysine. Toward this end, recombinant L71 containing Hyl was produced by co-expressing L71 and L230 in *E coli*. L71 was then isolated and glucosylated by purified recombinant L230 using UDP-Glc as the sugar donor. L71 containing glucosylhydroxylysine was extensively dialyzed before reacting with R699. R699-catalyzed glucosylation reaction was detected with a luciferase-based assay. However, no signal was found (data not shown). These findings do not support R699 functioning as a peptidyl glucosylhydroxylysine glucosyltransferase. Our work suggests that R699 acts on peptidyl galactosylhydroxylysine. The source of peptidyl galactosylhydroxylysine remains to be determined. It may be generated by the host or by an unknown mimiviral collagen hydroxylysyl galactosyltransferase. Since R655 shares moderate amino acid sequence identity (23%) with a human collagen hydroxylysyl galactosyltransferase GLT25D1, it warrants analysis of R655’s collagen hydroxylysyl galactosyltransferase activity.

**Figure 2.**
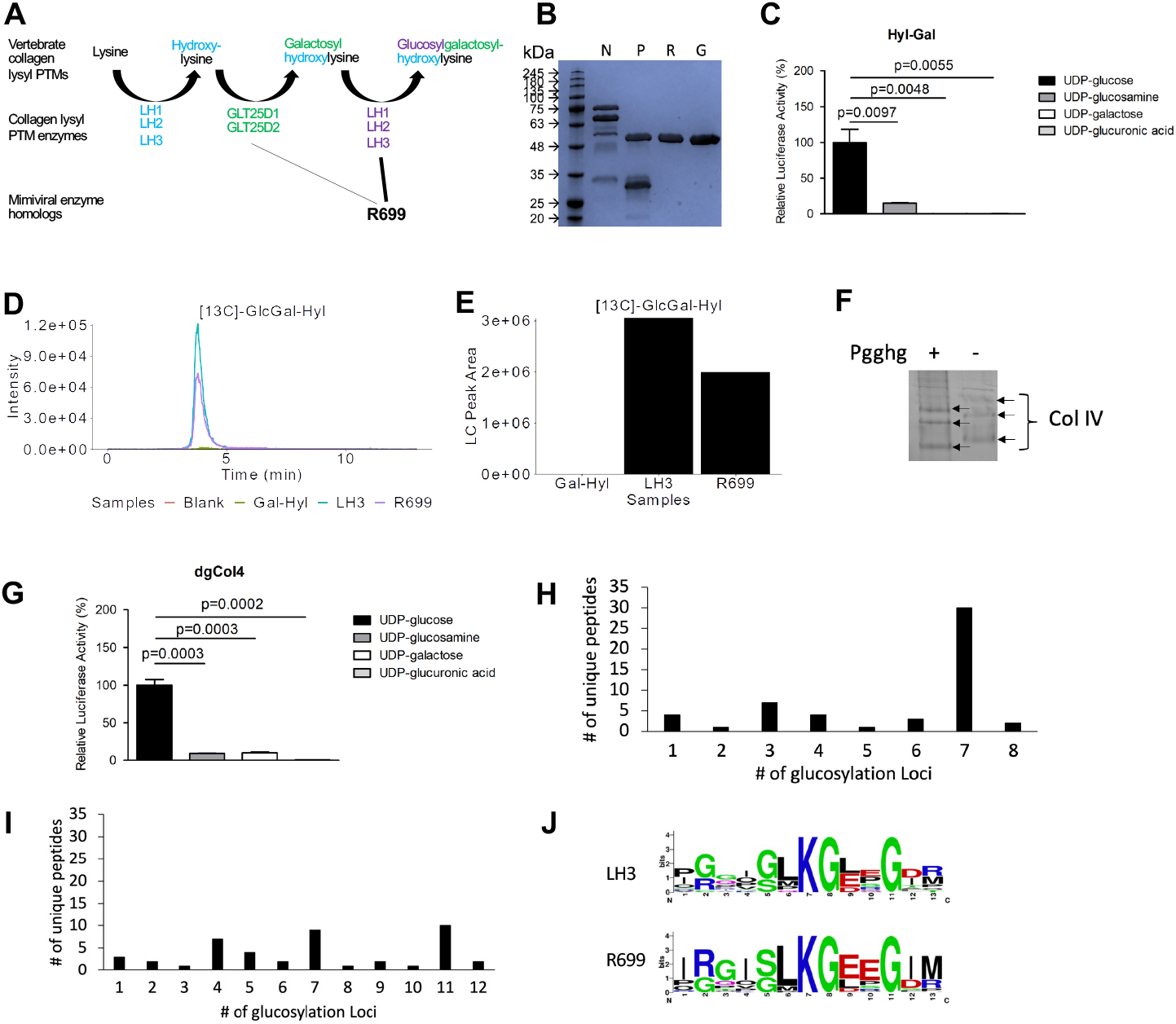
R699 is a collagen GGT. A, Schematics showing collagen hyl-*O*-linked glycosylation pathway. Sequence alignment shows that a mimiviral putative collagen glycosyltransferase R699 shares higher amino acid sequence identity (∼30%) with human collagen GGTs than hydroxylysyl galactosyltransferases. B, SDS-polyacrylamide gel electrophoresis of R699 protein after IMAC (N), PreScission cleavage (P), reverse IMAC (R) and gel filtration chromatography purification (G). R699 was purified close to homogeneity after 4-step purification. C, R699 GGT activity was assayed using an adenosine triphosphate-based luciferase assay. Substrate was hyl-gal and GGT activity was measured by detecting UDP production. *p* values, two-tailed Student’s *t* test. D and E, R699 GGT assay was performed using UDP-[UL-^13^C_6_] glucose. [^13^C]GlcGal-Hyl was confirmed with LC-MS analysis. The amount of [^13^C]GlcGal-Hyl was determined based on LC peaks using MultiQuant software (SCIEX). Chromatograms (D) and bar graph (E) were plotted using custom R scripts. F, Type IV collagen that had been pre-treated with wild-type (+) protein-glucosylgalactosylhydroxylysine glucosidase (PGGHG) or sham-treated was analyzed using SDS-polyacrylamide gel electrophoresis. G, R699 collagen GGT activity was assayed. Substrate was deglucosylated type IV collagen from F and GGT activity was measured similarly as in C. *p* values, two-tailed Student’s *t* test. H-J, R699 GGT assay was performed using deglucosylated type IV collagen and UDP-[UL-^13^C_6_] glucose. Peptides containing [^13^C]GlcGal-Hyl were detected by LC-MS (H). LH3 was assayed and detected as a positive control (I). The consensus motifs were generated using [^13^C]GlcGal-Hyl containing peptides (J).

### R699 protein, structure and its comparison with human collagen GGTs

To determine the structural basis of R699’s GGT activity, we solved the crystal structure of R699 with UDP-Glc (**Fig. 3A**). The structure was refined at a resolution of 2.39 Å with 2 molecules per asymmetric unit. The diffraction data statistics and refinement details were summarized in Supplementary Table. 12. The 2 R699 molecules in the asymmetric unit share high structural homology (root-mean-square deviation or RMSD = 0.2). Within each R699 molecule, two tandem domains with Rossmann fold are arranged similarly as in human GGT (44). The overall structure of catalytic and adjacent accessory domains shares moderate homology with human LH3 (PDB ID 6FXR) with an RMSD of 3.1. The catalytic domain of R699 shows higher structural similarity to the human LH3 GGT catalytic domain (PDB ID 6FXR) with an RMSD of 1.5, although they only share 34% sequence identity. As a comparison, a human LH3 GGT catalytic domain structure (PDB ID 6WFV) that we previously determined in a different crystallization condition superimposes with LH3 full length (PDB ID 6FXR) with an RMSD of 0.9. These findings suggest that R699 is a close structural homolog of human GGT.

**Figure 3.**
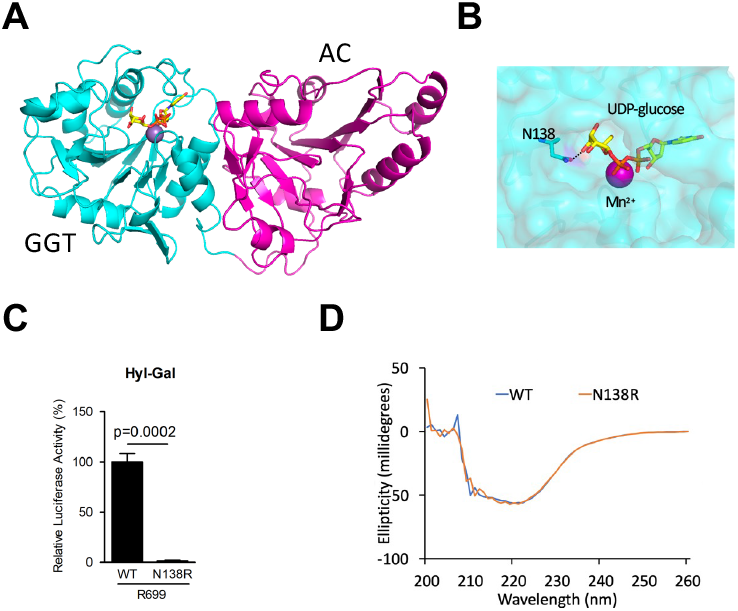
Structural insight into collagen GGT glucose binding. A, Ribbon diagram of the R699 GGT and accessory (AC) domains. Locations of Mn^2+^ (purple) and UDP-Glc (Yellow) are indicated. B, N138 OD1 atom of R699 interacts with the O3’ atom in the sugar moiety of UDP-Glc. C, N138R mutation abolishes R699 GGT activity. Substrate was hyl-gal and GGT activity was measured by detecting UDP production. *p* values, two-tailed Student’s *t* test. D, No alteration in R699 secondary structure due to loss of glucose binding residue. Circular dichroism spectrometry of wild-type R699 (WT) and R699 N138R mutant that is deficient in binding the sugar moiety of UDP-Glc. The proteins demonstrated similar spectra.

To the best of our knowledge, our structure is the first collagen GGT with a visible UDP-Glc bound. Interestingly, the sugar moiety resides in a previously unrecognized sugar-binding pocket with two glucose moieties in the asymmetric unit adopting similar conformations (**Fig. 3B** and supplemental Fig. 2). OD1 atom of Asn138 is involved in engaging glucose O3’ atom in both cases. By enzymatic activity assay, R699 GGT activity was lost upon N138R mutation (**Fig. 3C**), supporting a role of N138 in engaging glucose. To exclude the possibility that N138R mutation led to protein misfolding, we performed circular dichroism spectrometry and found no evidence for such misfolding (**Fig. 3D**). These findings validated our model that the sugar moiety of UDP-Glc resides in a previously unrecognized sugar-binding pocket, which are different from a recent work that suggested a glycoloop (Gly72-Gly87 in human LH3) was involved in sugar binding (45). The reason of inconsistency remains to be determined. It could be due to the different collagen GGT enzymes crystallized since the glycoloop is not conserved among different collagen GGTs (supplemental Fig. 3). It also could be due to different UDP-sugars utilized since the previous findings were based on UDP-xylose and UDP-glucuronic acid bound structures (45). In contrast, the genuine sugar donor UDP-Glc was utilized in the current study.

By comparing with the LH3 catalytic domain, we found that the R699 Mn^2+^-binding DXXD motif and UDP-binding Trp and Tyr residues are strictly conserved and positioned similarly in the structure (**Fig. 4A and 4E**). Site-directed mutagenesis and enzymatic activity assay showed that DXXD is critical for R699’s GGT activity (**Fig. 4B and 4C**). Interestingly, Trp but not Tyr is critical for GGT activity (**Fig. 4C**), suggesting Trp is the primary residue engaging UDP. Circular dichroism spectra suggest mutations are not deleterious to secondary structure (**Fig. 4D**). Asp190 and Asp191 in the poly-Asp repeat of LH3 that may be involved in catalysis are conserved in both identity and conformation (**Fig. 4E** arrows and supplemental Fig. 4). A unique Trp145 in LH3 that was thought to be a gating residue is also conserved but adopts a different conformation (**Fig. 4E** arrow and supplemental Fig. 4). Unlike the human GGT structure, the R699 inter-domain loop is fully visible and adopts a unique conformation (straight arrow in **Fig. 4F**). This loop is involved in R699 crystal packing contact, which may contribute to the conformation change observed. Sequence alignment suggested that the R699 inter-domain loop has a 5-residue deletion (**Fig. 4E** yellow shallow) and lacks the cysteine-linked hairpin structure (**Fig. 4E**). The largest 14-residue deletion (**Fig. 4E** brown shallow) happens in the accessory domain that is not required for LH3’s GGT activity but modulates LH2’s GGT activity (12,46). This largest deletion shortens 2 alpha helices and 1 beta strand, resulting in a major conformation change (curved arrow in **Fig. 4F**). The remaining deletions and an insertion that happen in the active and accessory domains have less impact on the overall structure.

**Figure 4.**
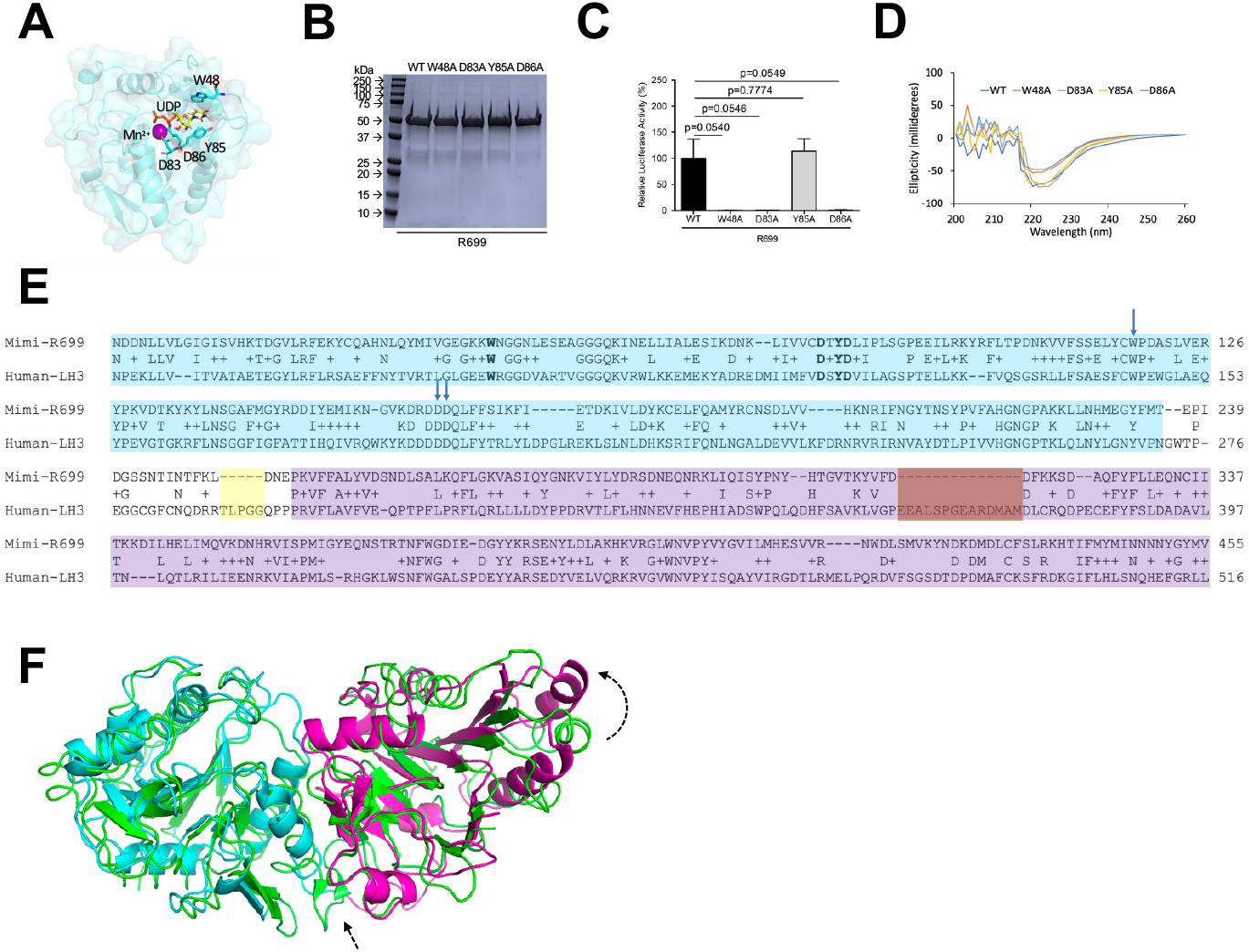
Key structural features are conserved among collagen GGTs. A, Mn^2+^ (purple ball) is coordinated by D83 and D86 while UDP (yellow, glucose moiety not shown) is sandwiched between W48 and Y85. B, SDS-polyacrylamide gel electrophoresis of R699 wild type (WT) and mutant recombinant proteins. C, GGT activity of WT and mutant R699 recombinant proteins was assayed using hyl-gal as substrate. The readout of the assay is adenosine triphosphate production which was detected using an adenosine triphosphate-based luciferase assay. *p* values, two-tailed Student’s *t* test. D, Circular dichroism spectrometry of wild-type R699 (WT) and R699 mutants that are Mn^2+^-and UDP-binding deficient. The proteins demonstrated similar spectra. E, Sequence alignment of R699 with human LH3. Residues within the GGT and AC domains are labeled in cyan and purple, respectively. Asp190 and Asp191 in poly-Asp repeat and Trp145 are indicated with arrows (residue number based on LH3 sequence). The interdomain loop deletion and the largest deletion in R699 is highlighted in yellow and brown squares, respectively. F, Superimposing R699 structure with human LH3 GGT and AC domains (PDB ID: 6FXR). The major differences between these two structures are indicated with dashed arrows.

### Website features and functionalities

To facilitate the further analysis of human and mimivirus homology, we established an interactive tool for easily searching and browsing of human and mimiviral homolog proteins (RRID Resource ID: SCR_022140 or https://guolab.shinyapps.io/app-mimivirus-publication/). Users can modify the search by changing the E value (Maximum Evalue), the length of query span in amino acid (Minimum Query Span) or percentage (Minimum QuerySpan Percent), the sequence identify percentage (Minimum Identity percentage) (**Fig. 5A**). The overall distribution of homologous proteins is shown in a histogram as counts vs. query length (**Fig. 5B**). If a list of all homologous proteins is needed, modify the search criteria without inputting a query. Clicking Excel on “Search Mimivirus Queries” tab or “Search Human Queries” tab will download an excel file including a list of all mimivirus or human homologous proteins, respectively (**Fig. 5C**). Search can be performed by inputting a query ID, gene name, or keywords about the query description. By clicking on the query_id, the details of the search (the Gene ID, symbol, description, and sequence) will be shown (**Fig. 5D**). The list of homologous hits is shown as a table under the Data table. The search details can be downloaded as an Excel file by clicking Excel (**Fig. 5D**).

**Figure 5.**
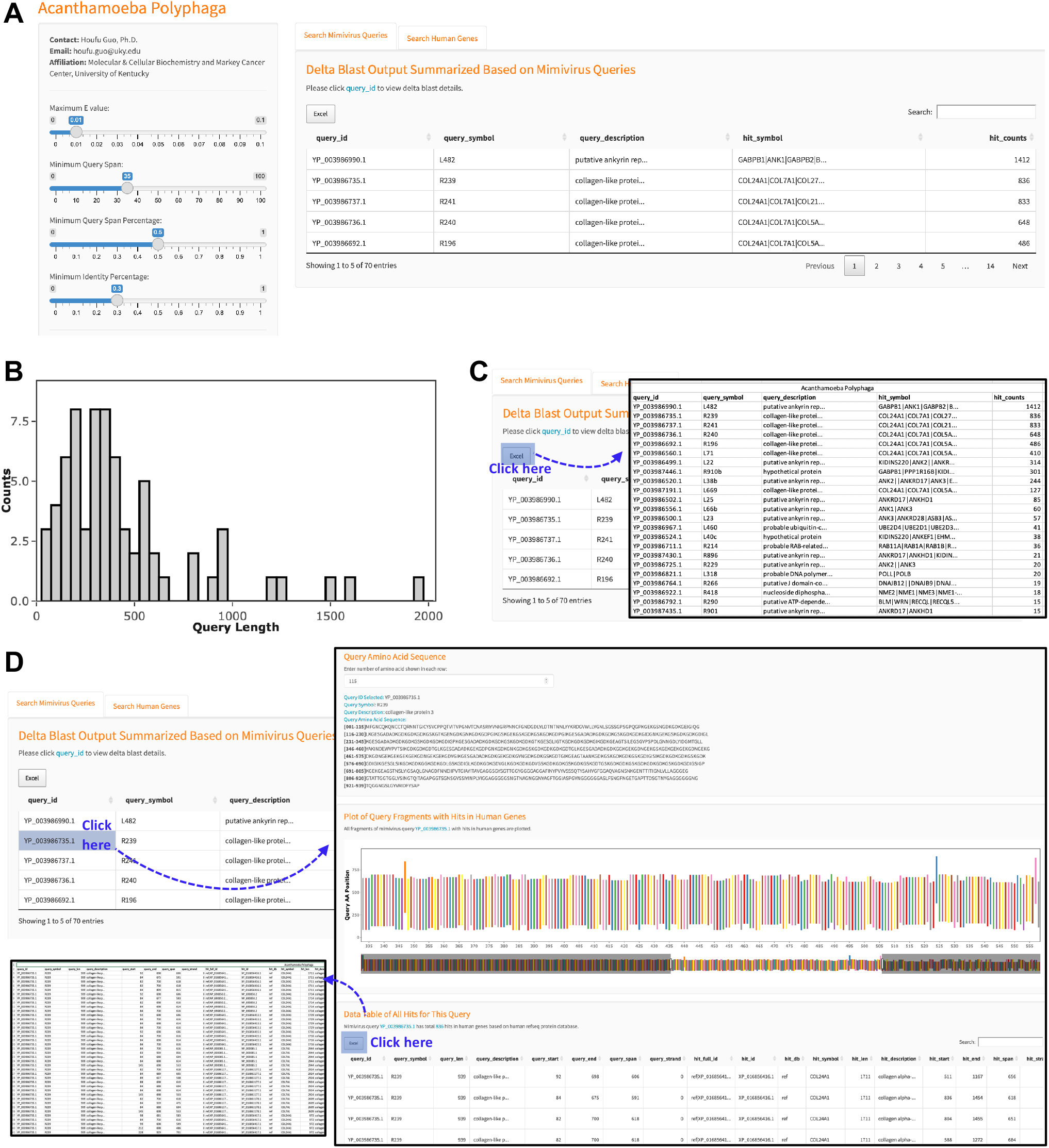
Human and mimivirus homology tool features and functionalities. A, An interactive tool was established for easily searching and browsing of human and mimiviral homolog proteins. B, Bar graph showing the overall distribution of homologous proteins. C, After performing search, clicking Excel (highlighted with a blue square) to download a list of human or mimivirus homologous proteins. D, Clicking on the query_id to show the details of the query and search (the Gene I, symbol, description and sequence). Under the section of Data Table of All Hits, an excel file about the detail of the search is available for download.

In conclusion, we performed genome-wide DELTA-BLAST of humans and mimivirus genomes and found 52 new mimiviral ORFs that show similarity to human proteins, including 8 mimiviral collagen-like proteins and 4 putative mimiviral collagen-modifying enzymes. Protein biochemistry and crystal structure studies confirmed the identify of a new collagen galactosylhydroxylysyl glucosyltransferase that engages UDP-glucose with a unique pocket, suggesting our search is robust. We established an interactive and searchable genome-wide comparison tool. This tool is established based on the DELTA-BLAST results that helped us identify more homologous proteins. The interactive and searchable nature of the website allows the users to modify the search criteria and quickly browse humans and mimivirus homologous proteins with different levels of homology at the genome-wide level.

## Experimental procedures

### Comparative genome-wide analysis of human and mimivirus homologous proteins

To search mimiviral protein sequences against the human non-redundant protein sequence database, we installed and ran the DELTA-BLAST command line application in Lipscomb Compute Cluster at the University of Kentucky with default parameters. We obtained Acanthamoeba Polyphaga Mimivirus GCF_000888735.1 assembly and annotation data from NCBI’s RefSeq ftp site ftp://ftp.ncbi.nlm.nih.gov/genomes/all/GCF/000/888/735/GCF_000888735.1_ViralProj60053. The file “*GCF_000888735.1_ViralProj60053_protein.faa.gz”* that contains 979 protein sequences was used as the input. All mimiviral protein sequences were searched against the human non-redundant protein sequence database, which was downloaded from NCBI’s blast ftp site https://ftp.ncbi.nlm.nih.gov/blast/db/. The file “*GCF_000888735.1_ViralProj60053_feature_table.txt.gz”* containing the feature information for all mimiviral protein sequences was used for annotation.

The DELTA-BLAST output in *xml* format was parsed and all high-scoring pairs (written as hits) were constructed into a tubular format using the biopython package *Bio. SearchIO*. The resulted data table was further processed in *R*. Among 979 mimiviral protein queries, 808 queries had at least one hit and 556603 hits were found in total. Using e value <= 0.01, hit span >= 35 amino acids, and the percentage of identical sequences between query and hit >= 0.25 as cutoffs, 356 query sequences with 85881 total hits passed the defined criteria. These 356 query sequences share sequence similarity with 4123 unique human RefSeq records in total.

To confirm the similarity mapping between mimivirus and human protein sequences, 4123 unique human RefSeq protein records we identified were DELTA-BLAST searched against the mimivirus non-redundant protein sequence database. The human RefSeq protein sequence file “GCF_000001405.39_GRCh38.p13_protein.faa.gz” of the latest GRCh38 assembly was downloaded from NCBI’s RefSeq ftp site ftp://ftp.ncbi.nlm.nih.gov/genomes/refseq/vertebrate_mammalian/Homo_sapiens/latest_assembly_versions/GCF_000001405.39_GRCh38.p13. Four RefSeq records were not present in the file therefore manually checked for their sequences in NCBI. A newly formed file containing all 4123 protein sequences of interest was used for DELTA-BLAST command line application with default parameters. After DELTA-BLAST, the output was processed in the same manner as the first DELTA-BLAST search. Using the same cutoffs, we found that 3325 human RefSeq records (1049 unique gene symbols) have hits in 307 unique mimiviral protein sequences. Of these 307 unique mimiviral protein sequences, 265 of them overlap with the 322 mimiviral queries that we identified in the first round of search. To summarize the results for overlapping mimiviral sequences, 1031 human genes and their corresponding mimiviral sequences were organized and presented (Supplementary Table. 13).

### Enrichment map and Network

To reduce the redundant hits between different databases, we selected the hits from the RefSeq database to perform pathway enrichment analysis using HUGO gene symbol, which resulted in 322 queries with at least one hit and 41520 hits in total. At the protein level, these 41520 hits are from 4123 unique RefSeq records, which contains 2027 proteins (IDs prefix with NP) and 2096 predicted proteins (IDs prefix with XP). These RefSeq record IDs were then converted into HUGO gene symbols using Bioconductor package *biomaRt*. The ones that cannot be converted by *biomaRt* were manually checked for the corresponding gene symbols in Genecard and BioGPS. Eventually, this conversion resulted in 1236 unique gene symbols, which were then used for pathway enrichment analysis and building Shiny App for visualization. The Shiny App is hosted on shinyapps.io server and is publicly available (https://guolab.shinyapps.io/app-mimivirus-publication/). This resource was submitted to RRID Portal with a Resource ID: SCR_022140.

To understand the overall biological and biochemical processes that the hits may be involved in, pathway enrichment analysis was performed using the R package *gprofiler2*. The significant GO and REACTOME pathways (adjusted p-value <= 0.05) with term sizes between 5 and 350 were selected for constructing pathway networks using *EnrichmentMap* (47). The resulted clusters were then automatically defined and summarized into major biological themes using *AutoAnnotate* (48). Finally, collagen-related pathways which formed the largest subnetwork were presented separately. All three steps were performed in *Cytoscape3.8.2* (49).

### Cloning, expression, and purification of R699 and variants

R699 gene was synthesized (Genscript). For enzymatic activity assay, R699 was cloned into a modified version of the pET28 vector using BamH1 and EcoR1 sites. This modified version of pET28 has PreScission and BamH1 recognition sites inserted to replace the thrombin recognition site. The endogenous BamH1 site was destroyed. Mutant constructs were generated using QuickChange Lightning Site-Directed Mutagenesis Kit (Agilent). For crystallization, R699 was cloned into a version of pET28-mCherry vector using BamH1 and EcoR1 sites. This pET28-mCherry vector has mCherry gene sequence and PreScission recognition site inserted between Nhe1 and BamH1 sites. All plasmids were verified by sanger sequencing and transformed into *E. coli* BL21 (NEB) for protein expression. Small scale R699-BL21 overnight culture with 50 mg per liter of kanamycin (GoldBio) was prepared and 10 ml of small-scale overnight culture was used to inoculate 800 ml large scale culture using Terrific Broth Medium (Alpha Biosciences) in the presence of the same amount of kanamycin. Culture was grown at 37°C to OD_600_= 1.5, induced with 1 mM isopropyl β-D-1-thiogalactopyranoside (IPTG, GoldBio) and grown at 16°C for 18 hours. Cells were collected, pelleted and then resuspended in binding buffer (20 mM Tris, pH 8.0, 200 mM NaCl and 15 mM imidazole). The cells were lysed by sonication and then centrifuged at 23,000g for 15 min. The recombinant R699 proteins (wild type or mutants) were purified with immobilized metal affinity chromatography and eluted with elution buffer (200 mM NaCl and 300 mM imidazole, pH 8.0). For enzymatic activity assay, R699 protein was dialyzed at 16°C for 18 hours in 20 mM HEPES, pH 7.4, 150 mM NaCl.

### Crystallization, structure determination, and refinement

mCherry-R699 was first purified with immobilized metal affinity chromatography as described above. The eluted recombinant protein was cleaved with PreScission protease at 4°C for 18 hours while dialyzing in gel filtration buffer (20 mM Tris, pH 8.0, 200 mM NaCl). After PreScission protease cleavage, R699 was purified with reverse immobilized metal affinity chromatography to remove mCherry protein and other contaminants that bind to nickel resin. The eluted protein was further separated by gel filtration using a Hiload 16/60 Superdex 200 PG column at a flow rate of 1 ml per minute. Peak fractions were combined and concentrated for crystal trials. Single, high-quality crystals with two molecules in the asymmetric unit were obtained via hanging drop vapor diffusion using a Mosquito liquid handling robot (TTPLabtech) using a 200-nL drop. R699 (16 mg mL^-1^) supplemented with 10 mM uracil-diphosphate glucose and 2 mM manganese(II) chloride was mixed with 200 mM NaCl, 0.1M Na/K phosphate pH 6.2 and 40% (v v^-1^) PEG 400 at 1:1 ratio and incubated at 18°C. Diffraction data were collected on the 22-ID beamline of SERCAT at the Advanced Photon Source, Argonne National Laboratory (Supplementary Table. 12) at 110K at a wavelength of 1.0 Å. Data were processed using CCP4, version 7.1.018 (50) and the structure was solved by molecular replacement with Phenix using RoseTTAFold and AlphaFold models as search templates (51-53). The structure was then fully built and refined via iterative model building and refinement using Coot and Phenix, respectively (53,54). Protein structure similarity was compared using the Dali server (55). Structure interface was analyzed using protein interfaces, surfaces and assemblies’ service PISA at the European Bioinformatics Institute (56). Molecular graphics were prepared using Pymol (DeLano, W. L. The PyMOL molecular graphics system. http://www.pymol.org). Amino acid sequence alignment of R699 and human GGTs was performed using Clustal Omega (57).

### GGT enzymatic activity assay

GGT activity was measured similarly as previously described (12). The assay was performed in reaction buffer (100 mM HEPES buffer pH 8.0, 150 mM NaCl) at 37 °C for 1 h with 1 μM R699 enzyme, 100 μM MnCl_2_, 200 μM UDP-Glc (MilliporeSigma, St. Louis, MO), 1 mM dithiothreitol and 1.75 mM galactosyl hydroxylysine (Gal-Hyl, Cayman Chemical, Ann Arbor, MI) or 2 μM deglucosylated collagen IV. Deglucosylated collagen IV was generated using a glycosidase PGGHG as previously described (12). GGT activity was measured by detecting UDP production with an ATP–based luciferase assay (UDP-Glo™ Glycosyltransferase Assay, Promega, Madison, WI) according to manufacturers’ instructions. Experiments were performed in triplicate from distinct samples, and an unpaired t-test was used to compare the enzymatic activity of different samples. The glucosylation of galactosyl hydroxylysine was further confirmed by mass spectrometry.

### Mass spectrometry

To confirm the glucosylation of Gal-Hyl and type IV collagen by R699, the R699 GGT assay was performed similarly as discussed above, except that UDP-Glc was replaced with the same concentration of UDP-[UL-^13^C_6_] glucose (Omicron Biochemicals, Inc). LC-MS analysis was used to detect [^13^C]GlcGal-Hyl. LH3 catalyzed GGT activity assay was used as a positive control. For LC-MS analysis of [^13^C]GlcGal-Hyl, R699 and LH3 assay samples were diluted to ∼1 μM in 50% acetonitrile containing 0.1% formic acid. LC-MS analysis was performed using a 1260 Infinity UHPLC System (Agilent) coupled to a Qtrap 6500 mass spectrometer (SCIEX). Samples were separated on a Kinetex EVO C18 column (Phenomenex) with mobile phases included: A) water + 0.1% formic acid, B) acetonitrile + 0.1% formic acid. LC peaks were integrated using MultiQuant software (SCIEX). Peak areas and chromatograms were plotted using custom R scripts. Experiments were performed once.

For mapping the glucosylation loci, 25ug of type IV collagen and collagen GGTs from each GGT reaction mixture was solubilized with 5% SDS, 50mM TEAB, pH 7.55, final volume 25uL. The sample was then centrifuged at 17,000g for 10 minutes to remove any debris. Proteins were reduced by making the solution 20mM TCEP (Thermo, #77720) and incubated at 65°C for 30min. The sample was cooled to RT and 1 uL of 0.5M iodoacetamide acid added and allowed to react for 20 minutes in the dark. 2.75 ul of 12% phosphoric acid is added to the protein solution. 165uL of binding buffer (90% Methanol, 100mM TEAB final; pH 7.1) is then added to the solution. The resulting solution is added to S-Trap spin column (protifi.com) and passed through the column using a bench top centrifuge (30s spin at 4,000g). The spin column is washed with 400uL of binding buffer and centrifuged. This is repeated two more times. Trypsin is added to the protein mixture in a ratio of 1:25 in 50mM TEAB, pH=8, and incubated at 37°C for 4 hours. Peptides were eluted with 80uL of 50mM TEAB, followed by 80uL of 0.2% formic acid, and finally 80 uL of 50% acetonitrile, 0.2% formic acid. The combined peptide solution is then dried in a speed vac and resuspended in 2% acetonitrile, 0.1% formic acid, 97.9% water and placed in an autosampler vial.

Peptide mixtures were analyzed by nanoflow liquid chromatography-tandem mass spectrometry (nanoLC-MS/MS) using a nano-LC chromatography system (UltiMate 3000 RSLCnano, Dionex), coupled on-line to a Thermo Orbitrap Fusion mass spectrometer (Thermo Fisher Scientific, San Jose, CA) through a nanospray ion source (Thermo Scientific). A trap and elute method was used. The trap column was a C18 PepMap100 (300um X 5mm, 5um particle size) from ThermoScientific. The analytical columns was an Acclaim PepMap 100 (75um X 25 cm) from (Thermo Scientific). Peptides were eluted using a 120 min gradient (mobile phase A = 0.1% formic acid (Thermo Fisher), mobile phase B = 99.9% acetonitrile with 0.1% formic acid (Thermo Fisher); hold 12% B for 5 min, 2-6% B in 0.1 min, 6-25% in 100 min, 25-50% in 15 min) at a flow rate of 350 nL min^-1^. Eluted peptide ions were analyzed using a data-dependent acquisition (DDA) method with resolution settings of 120,000 and 15,000 (at *m/z* 200) for MS1 and MS2 scans, respectively. DDA-selected peptides were fragmented using stepped high energy collisional dissociation (27, 32, 37%).

Tandem mass spectra were extracted and charge state deconvoluted by Proteome Discoverer (Thermo Fisher, version 1.4.1.14). Deisotoping is not performed. All MS/MS spectra were searched against a Uniprot Ecoli and human databases as background, and a custom database made of common contaminants and collagen proteins using Sequest and MS Amanda search engines. Searches were performed with a parent ion tolerance of 10 ppm and a fragment ion tolerance of 0.02 Da. Trypsin is specified as the enzyme, allowing for two missed cleavages. Fixed modification of carbamidomethyl (C) and variable modifications of oxidation (M, K, and P), ^13^C_6_-glucosylgalactosyl (+346.121 Da (K)), Galactosyl +178.048 Da (K), and Glucosylgalactosyl +340.101 Da (K). Consensus motifs were generated using WebLogo (58).

### Circular dichroism

Circular dichroism spectra were measured using a J-810 spectrapolarimeter (Jasco, Easton, MD) with a 2 mm path length quartz cuvette. All measurements were performed at 20 °C. Three scans were averaged to generate each spectrum. A blank spectrum of buffer was collected in the same manner and used for background subtraction. For **Fig. 3E**, R699 wild type and mutant recombinant proteins were dialyzed and measured in 20 mM HEPES and 150 mM NaCl (pH 7.4). For **Fig. 4D**, R699 recombinant proteins were analyzed in 0.01 M sodium phosphate, 150 mM NaCl (pH 7.4) and 10% glycerol at a concentration of 0.5 mg ml^−1^. Results represent the mean values from triplicate technical repeats in a single experiment. Each protein was analyzed once.

## Data availability

Crystal structure has been deposited in the Worldwide Protein Data Bank under RCSB accession ID number 7UL9.

## Supporting information

This article contains supporting information.

## Acknowledgments

We thank Drs. Jonathan Kurie from MD Anderson Cancer Center, Emilia Galperin, Trevor Creamer, Louis B. Hersh, Craig Vander Kooi and Matthew Gentry from the University of Kentucky for sharing equipment, reagents, and helpful discussions. We thank Dr. Trevor Romsdahl for assistance with the LCMS analysis.

## Funding and additional information

This work was supported by the National Institutes of Health grant R00CA225633 (H.G.). WKR and the UTMB Mass Spectrometry Facility receives support from Cancer Prevention Research Institute of Texas (CPRIT) grant number RP190682. The content is solely the responsibility of the authors and does not necessarily represent the official views of the National Institutes of Health.

## Conflict of interest

The authors declare that they have no conflicts of interest with the contents of this article.

## Notes

https://guolab.shinyapps.io/app-mimiviruspublication/

### Competing Interest Statement

The authors have declared no competing interest.

