## Supplemental figures for "Comparative genomic and crystal structure analyses identify a collagen glucosyltransferase from *Acanthamoeba Polyphaga Mimivirus*"

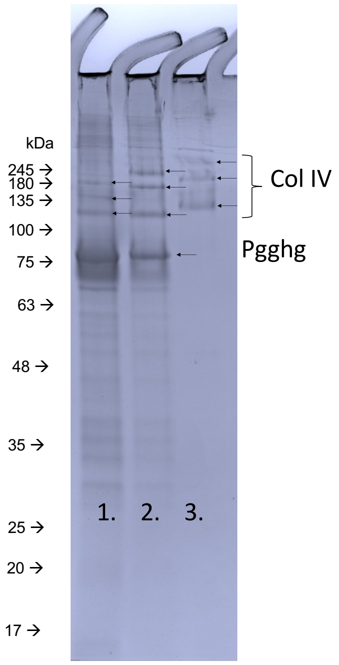


1. Denatured collagen IV + high concentration PGGHG
2. Denatured collagen IV + low concentration PGGHG
3. Denatured collagen IV

**Figure S1. Uncut SDS-polyacrylamide gel for Fig. 2F.** Type IV collagen that had been pre-treated with wild-type (+) protein glucosylgalactosylhydroxylysine glucosidase (PGGHG) or sham-treated was analyzed using SDS-polyacrylamide gel electrophoresis.

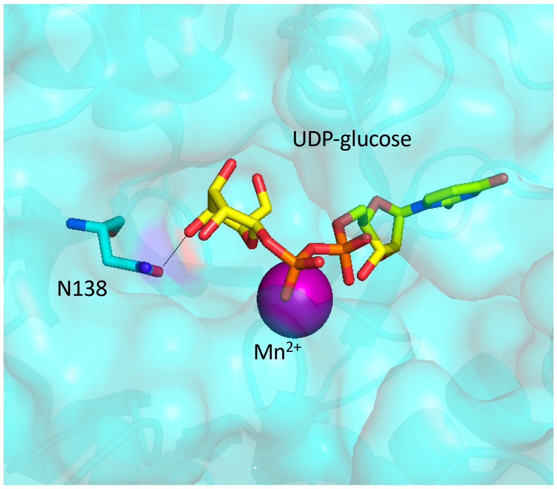


**Figure S2. R699 binds UDP-Glc similarly in the other subunit with N138 OD1 atom of R699 engaging O3' atom in the sugar moiety of UDP-glucose.**

```
R699        -----------------------------GEQSNNDDNLLVLGIGISVHKTDGVLRFEKY      31
hLH3        MTSSGPGPRFLLLLPLLLPPAASASDRPRGRDPVNPEKLLVI--TVATAETEGYLRFLRS      58
hLH1        -----MRPLLL---LALLGWLLL---AEAKGDAKPEDNLLVL--TVATKETEGFRRFKRS      47
hLH2        MGGCTVKPQLLLLALVLHPWNPCLGADSEKPSSIPTDKLLVI--TVATKESDGFHRFMQS      58
                                           .    ::***:   ::. :::*  ** :

R699        CQAHNLQYMIVGEGKKWNGGNLESEAGGGQKINELLIALESIKD--NKLIVVCDTYDLIP      89
hLH3        AEFFNYTVRTLGLGEEWRGGDVARTVGGGQKVRWLKKEMEKYADREDMIIMFVDSYDVIL      118
hLH1        AQFFNYKIQALGLGEDWNVEKGT-SAGGGQKVRLLKKALEKHADKEDLVILFTDSYDVLF      106
hLH2        AKYFNYTVKVLGQGEEWRGGDGINSIGGGQKVRLMKEVMEHYADQDDLVVMFTECFDVIF      118
            .: .*     :* *:.*.  .     *****:. :   :*   *  : :::. : :*::

R699        LSGPEEILRKYRFLTPDNKVVFSSELYCWPDASLVERYPKVDTKYKYLNSGAFMGYRDDI      149
hLH3        AGSPTELLKKFV--QSGSRLLFSAESFCWPEWGLAEQYPEVGTGKRFLNSGGFIGFATTI      176
hLH1        ASGPRELLKKFR--QSRSQVVFSAEELIYPDRRLETKYPVVSDGKRFLGSGGFIGYAPNL      164
hLH2        AGGPEEVLKKFQ--KANHKVVFAADGILWPDKRLADKYPVVHIGKRYLNSGGFIGYAPYV      176
             ..* *:*:*:       :::*:::   :*:  *  :** *    ::*.**.*:*:   :

R699        YEMIKN-GVKDRDDDQLFFSIKFIETD-----KIVLDYKCELFQAMYRCNSDLVVH----      199
hLH3        HQIVRQWKYKDDDDDQLFYTRLYLDPGLREKLSLNLDHKSRIFQNLSGALDEVVLKFDRN      236
hLH1        SKLVAEWEGQDSDSDQLFYTKIFLDPEKREQINITLDHRCRIFQNLDGALDEVVLKFEMG      224
hLH2        NRIVQQWNLQDNDDDQLFYTKVYIDPLKREAINITLDHKCKIFQTLNGAVDEVVLKFENG      236
             .:: :   :* *.****::  :::       .: **::..:** :  . .::*::

R699        KNRIFNGYTNSYPVFAHGNGPAKKLLNHMEGYFMTEPIDGSSN--------TINTFKLDN      251
hLH3        RVRIRNVAYDTLPIVVHGNGPTKLQLNYLGNYVPNGWTPEGGCGFCNQDRRTLPGG--QP      294
hLH1        HVRARNLAYDTLPVLIHGNGPTKLQLNYLGNYIPRFWTFETGCTVCDEGLRSLKGIGDEA      284
hLH2        KARAKNTFYETLPVAINGNGPTKILLNYFGNYVPNSWTQDNGCTLCEFDTVDLSAV--DV      294
            : *  *   :: *:  :****:*  **:: .*.        .          :     :

R699        EPKVFFALYVDSNDLSALKQFLGKVASIQYGNKVIYLYDRSDNEQNRKLIQISYPNYHT-      310
hLH3        PPRVFLAVFVEQPT-PFLPRFLQRLLLLDYPPDRVTLFLHNNEVFHEPHIADSWPQLQDH      353
hLH1        LPTVLVGVFIEQPT-PFVSLFFQRLLRLHYPQKHMRLFIHNHEQHHKAQVEEFLAQHGSE      343
hLH2        HPNVSIGVFIEQPT-PFLPRFLDILLTLDYPKEALKLFIHNKEVYHEKDIKVFFDKAKHE      353
             * * ..::::.     :  *:  :  :.*  . : *: :..:  :.  :     :

R699        ---------------GVTKYVFDDFK--KSDAQFYFLLEQNCIITKKDILHELIMQVKDN      353
hLH3        FSAVKLVGPEEALSPGEARDMAMDLCRQDPECEFYFSLDADAVLTNLQTLRIL---IEEN      410
hLH1        YQSVKLVGPEVRMANADARNMGADLCRQDRSCTYYFSVDADVALTEPNSLRLL---IQQN      400
hLH2        IKTIKIVGPEENLSQAEARNMGMDFCRQDEKCDYYFSVDADVVLTNPRTLKIL---IEQN      410
                           . :: :  *:   . .. :** :: :  :*:   *: *   :::*

R699        HRVISPMIGYEQNSTRTNFWGDI-EDGYYKRSENYLDLAKHKVRGLWNVPYVYGVILMHE      412
hLH3        RKVIAPMLSR-HGKLWSNFWGALSPDEYYARSEDYVELVQRKRVGVWNVPYISQAYVIRG      469
hLH1        KNVIAPLMTR-HGRLWSNFWGALSADGYYARSEDYVDIVQGRRVGVWNVPYISNIYLIKG      459
hLH2        RKIIAPLVTR-HGKLWSNFWGALSPDGYYARSEDYVDIVQGNRVGVWNVPYMANVYLIKG      469
            :.:*:*::   :.   :**** :  * **  ***:*:::.: .  *:*****:    :::

R699        SVVRN----WDLSMVKYNDKDMDLCFSLRKHTIFMYMINNNNYGYMV------------      455
hLH3        DTLRMELPQRDVFSGSDTDPDMAFCKSFRDKGIFLHLSNQHEFGRLLATSRYDTEHLHPD      529
hLH1        SALRGELQSSDLFHHSKLDPDMAFCANIRQQDVFMFLTNRHTLGHLLSLDSYRTTHLHND      519
hLH2        KTLRSEMNERNYFVRDKLDPDMALCRNAREMGVFMYISNRHEFGRLLSTANYNTSHYNND      529
            ..:*      :    .  * ** :* . *.   :*:.: *.:  * ::
```

**Figure S3. Sequence alignment of R699 with human LH1-3.** Residues within the previously proposed glycoloop are labeled in a blue square.

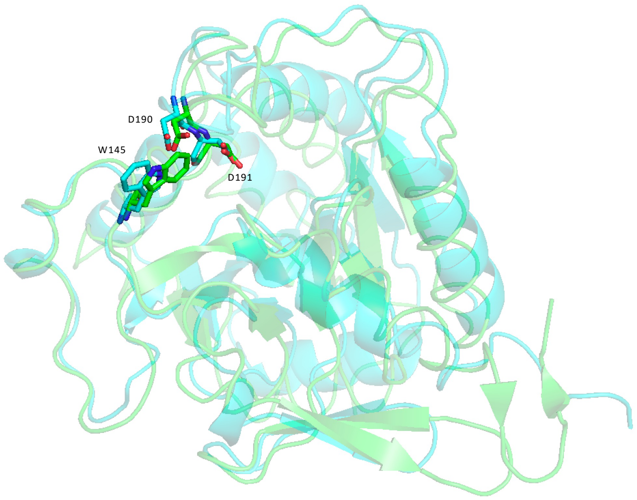


**Figure S4. Ribbon diagram of the R699 (cyan) and LH3 (green) GGT domain.** The conformation of D190, D191 and W145 in LH3 in comparison to the corresponding residues D162, D163 and W118 in R699.
