## Supplemental Table 5 for "Comparative genomic and crystal structure analyses identify a collagen glucosyltransferase from *Acanthamoeba Polyphaga Mimivirus*"

| id | source | term_id | term_name | term_size | intersection_size | | p_value |
|---|---|---|---|---|---|---|---|
| 1 | GO:BP | GO:0000209 | protein polyubiquitination | 327 | 88 | | 5.1e−27 |
| 2 | GO:BP | GO:0006457 | protein folding | 195 | 49 | | 1.0e−12 |
| 3 | GO:BP | GO:0051085 | chaperone cofactor−dependent protein refolding | 24 | 17 | | 6.1e−12 |
| 4 | GO:BP | GO:0031146 | SCF−dependent proteasomal ubiquitin−dependent protein catabolic process | 93 | 32 | | 8.9e−12 |
| 5 | GO:BP | GO:0051084 | 'de novo' posttranslational protein folding | 29 | 17 | | 6.6e−10 |
| 6 | GO:BP | GO:0006458 | 'de novo' protein folding | 33 | 18 | | 7.2e−10 |
| 7 | GO:BP | GO:0061077 | chaperone−mediated protein folding | 53 | 22 | | 2.2e−09 |
| 8 | GO:BP | GO:0070936 | protein K48−linked ubiquitination | 55 | 22 | | 5.4e−09 |
| 9 | GO:BP | GO:0030198 | extracellular matrix organization | 320 | 55 | | 2.9e−07 |
| 10 | GO:BP | GO:0043062 | extracellular structure organization | 321 | 55 | | 3.2e−07 |
| 11 | GO:BP | GO:0048524 | positive regulation of viral process | 100 | 26 | | 3.8e−06 |
| 12 | GO:BP | GO:0006260 | DNA replication | 227 | 42 | | 4.7e−06 |
| 13 | GO:BP | GO:0035966 | response to topologically incorrect protein | 177 | 36 | | 4.9e−06 |
| 14 | GO:BP | GO:0046777 | protein autophosphorylation | 209 | 39 | | 1.4e−05 |
| 15 | GO:BP | GO:0042026 | protein refolding | 19 | 11 | | 1.4e−05 |
| 16 | GO:BP | GO:0048705 | skeletal system morphogenesis | 88 | 23 | | 2.9e−05 |
| 17 | GO:BP | GO:0090084 | negative regulation of inclusion body assembly | 10 | 8 | | 4.1e−05 |
| 18 | GO:BP | GO:0001501 | skeletal system development | 281 | 46 | | 4.6e−05 |
| 19 | GO:BP | GO:0000086 | G2/M transition of mitotic cell cycle | 238 | 41 | | 6.2e−05 |
| 20 | GO:BP | GO:0044839 | cell cycle G2/M phase transition | 257 | 43 | | 6.9e−05 |
| 21 | GO:BP | GO:0048704 | embryonic skeletal system morphogenesis | 37 | 14 | | 1.3e−04 |
| 22 | GO:BP | GO:0009952 | anterior/posterior pattern specification | 95 | 23 | | 1.4e−04 |
| 23 | GO:BP | GO:0006986 | response to unfolded protein | 155 | 30 | | 3.4e−04 |
| 24 | GO:BP | GO:0017157 | regulation of exocytosis | 147 | 29 | | 3.6e−04 |
| 25 | GO:BP | GO:0070979 | protein K11−linked ubiquitination | 29 | 12 | | 3.6e−04 |
| 26 | GO:BP | GO:0006301 | postreplication repair | 52 | 16 | | 4.2e−04 |
| 27 | GO:BP | GO:0090083 | regulation of inclusion body assembly | 13 | 8 | | 9.7e−04 |
| 28 | GO:BP | GO:0046782 | regulation of viral transcription | 62 | 17 | | 1.1e−03 |
| 29 | GO:BP | GO:0006904 | vesicle docking involved in exocytosis | 32 | 12 | | 1.3e−03 |
| 30 | GO:BP | GO:0018105 | peptidyl−serine phosphorylation | 266 | 41 | | 1.4e−03 |
| 31 | GO:BP | GO:0050792 | regulation of viral process | 210 | 35 | | 1.5e−03 |
| 32 | GO:BP | GO:0048706 | embryonic skeletal system development | 50 | 15 | | 1.5e−03 |
| 33 | GO:BP | GO:0043903 | regulation of symbiotic process | 222 | 36 | | 2.0e−03 |
| 34 | GO:BP | GO:0006261 | DNA−dependent DNA replication | 142 | 27 | | 2.0e−03 |
| 35 | GO:BP | GO:0009262 | deoxyribonucleotide metabolic process | 28 | 11 | | 2.3e−03 |
| 36 | GO:BP | GO:0006289 | nucleotide−excision repair | 105 | 22 | | 3.9e−03 |
| 37 | GO:BP | GO:0006354 | DNA−templated transcription, elongation | 106 | 22 | | 4.6e−03 |
| 38 | GO:BP | GO:0034976 | response to endoplasmic reticulum stress | 269 | 40 | | 4.9e−03 |
| 39 | GO:BP | GO:0003002 | regionalization | 158 | 28 | | 5.6e−03 |
| 40 | GO:BP | GO:0018126 | protein hydroxylation | 25 | 10 | | 5.9e−03 |
| 41 | GO:BP | GO:0030433 | ubiquitin−dependent ERAD pathway | 77 | 18 | | 6.5e−03 |
| 42 | GO:BP | GO:0018209 | peptidyl−serine modification | 284 | 41 | | 7.9e−03 |
| 43 | GO:BP | GO:0071897 | DNA biosynthetic process | 152 | 27 | | 8.0e−03 |
| 44 | GO:BP | GO:0009408 | response to heat | 128 | 24 | | 1.1e−02 |
| 45 | GO:BP | GO:0006283 | transcription−coupled nucleotide−excision repair | 72 | 17 | | 1.1e−02 |
| 46 | GO:BP | GO:1904355 | positive regulation of telomere capping | 17 | 8 | | 1.4e−02 |
| 47 | GO:BP | GO:0050434 | positive regulation of viral transcription | 39 | 12 | | 1.5e−02 |
| 48 | GO:BP | GO:0085020 | protein K6−linked ubiquitination | 9 | 6 | | 1.6e−02 |
| 49 | GO:BP | GO:0036503 | ERAD pathway | 99 | 20 | | 2.0e−02 |
| 50 | GO:BP | GO:0048598 | embryonic morphogenesis | 307 | 42 | | 2.4e−02 |
| 51 | GO:BP | GO:0030199 | collagen fibril organization | 41 | 12 | | 2.6e−02 |
| 52 | GO:BP | GO:0035967 | cellular response to topologically incorrect protein | 155 | 26 | | 3.5e−02 |
| 53 | GO:BP | GO:0034605 | cellular response to heat | 111 | 21 | | 3.6e−02 |
| 54 | GO:BP | GO:0070841 | inclusion body assembly | 19 | 8 | | 3.9e−02 |
| 55 | GO:BP | GO:0009266 | response to temperature stimulus | 156 | 26 | | 4.0e−02 |
| 56 | GO:BP | GO:0006368 | transcription elongation from RNA polymerase II promoter | 79 | 17 | | 4.0e−02 |
| 57 | GO:BP | GO:1900034 | regulation of cellular response to heat | 79 | 17 | | 4.0e−02 |
| 58 | GO:BP | GO:1901796 | regulation of signal transduction by p53 class mediator | 167 | 27 | | 4.8e−02 |

g:Profiler (biit.cs.ut.ee/gprofiler)
