## Supplemental Table 6 for "Comparative genomic and crystal structure analyses identify a collagen glucosyltransferase from *Acanthamoeba Polyphaga Mimivirus*"

| id | source | term_id | term_name | term_size | intersection_size | | p_value |
|---|---|---|---|---|---|---|---|
| 1 | GO:CC | GO:0005581 | collagen trimer | 32 | 29 | | 4.6e−29 |
| 2 | GO:CC | GO:0000151 | ubiquitin ligase complex | 275 | 79 | | 8.1e−28 |
| 3 | GO:CC | GO:0098644 | complex of collagen trimers | 20 | 19 | | 1.8e−19 |
| 4 | GO:CC | GO:0031461 | cullin−RING ubiquitin ligase complex | 161 | 50 | | 9.6e−19 |
| 5 | GO:CC | GO:0005788 | endoplasmic reticulum lumen | 282 | 64 | | 2.5e−16 |
| 6 | GO:CC | GO:0019005 | SCF ubiquitin ligase complex | 60 | 28 | | 2.4e−15 |
| 7 | GO:CC | GO:0031463 | Cul3−RING ubiquitin ligase complex | 36 | 18 | | 4.2e−10 |
| 8 | GO:CC | GO:0098643 | banded collagen fibril | 11 | 10 | | 4.8e−09 |
| 9 | GO:CC | GO:0005583 | fibrillar collagen trimer | 11 | 10 | | 4.8e−09 |
| 10 | GO:CC | GO:0098651 | basement membrane collagen trimer | 9 | 9 | | 7.1e−09 |
| 11 | GO:CC | GO:0098645 | collagen network | 8 | 8 | | 1.1e−07 |
| 12 | GO:CC | GO:0098642 | network−forming collagen trimer | 8 | 8 | | 1.1e−07 |
| 13 | GO:CC | GO:0005587 | collagen type IV trimer | 7 | 7 | | 1.7e−06 |
| 14 | GO:CC | GO:0005604 | basement membrane | 59 | 17 | | 4.4e−05 |
| 15 | GO:CC | GO:0005663 | DNA replication factor C complex | 5 | 5 | | 3.9e−04 |
| 16 | GO:CC | GO:0008540 | proteasome regulatory particle, base subcomplex | 12 | 7 | | 9.9e−04 |
| 17 | GO:CC | GO:0032593 | insulin−responsive compartment | 9 | 6 | | 1.8e−03 |
| 18 | GO:CC | GO:0030139 | endocytic vesicle | 254 | 36 | | 3.3e−03 |
| 19 | GO:CC | GO:0005657 | replication fork | 62 | 14 | | 1.2e−02 |
| 20 | GO:CC | GO:0055037 | recycling endosome | 140 | 23 | | 1.3e−02 |
| 21 | GO:CC | GO:0045335 | phagocytic vesicle | 98 | 18 | | 2.0e−02 |
| 22 | GO:CC | GO:0034663 | endoplasmic reticulum chaperone complex | 5 | 4 | | 2.8e−02 |
| 23 | GO:CC | GO:0031371 | ubiquitin conjugating enzyme complex | 9 | 5 | | 4.0e−02 |
| 24 | GO:CC | GO:0016607 | nuclear speck | 320 | 39 | | 4.5e−02 |
| 25 | GO:CC | GO:0005665 | RNA polymerase II, core complex | 14 | 6 | | 4.9e−02 |

g:Profiler (biit.cs.ut.ee/gprofiler)
