## Supplemental Table 7 for "Comparative genomic and crystal structure analyses identify a collagen glucosyltransferase from *Acanthamoeba Polyphaga Mimivirus*"

| id | source | term_id | term_name | term_size | intersection_size | | p_value |
|---|---|---|---|---|---|---|---|
| 1 | GO:MF | GO:0030020 | extracellular matrix structural constituent conferring tensile strength | 41 | 39 | | 3.2e−41 |
| 2 | GO:MF | GO:0005201 | extracellular matrix structural constituent | 167 | 66 | | 1.1e−31 |
| 3 | GO:MF | GO:0016887 | ATPase activity | 316 | 85 | | 6.1e−27 |
| 4 | GO:MF | GO:0061650 | ubiquitin−like protein conjugating enzyme activity | 42 | 32 | | 7.0e−27 |
| 5 | GO:MF | GO:0061631 | ubiquitin conjugating enzyme activity | 40 | 31 | | 2.1e−26 |
| 6 | GO:MF | GO:0004386 | helicase activity | 101 | 39 | | 1.1e−17 |
| 7 | GO:MF | GO:0019003 | GDP binding | 66 | 30 | | 1.1e−15 |
| 8 | GO:MF | GO:0004707 | MAP kinase activity | 20 | 17 | | 4.8e−15 |
| 9 | GO:MF | GO:0005524 | ATP binding | 285 | 63 | | 8.1e−15 |
| 10 | GO:MF | GO:0004693 | cyclin−dependent protein serine/threonine kinase activity | 23 | 18 | | 8.2e−15 |
| 11 | GO:MF | GO:0097472 | cyclin−dependent protein kinase activity | 23 | 18 | | 8.2e−15 |
| 12 | GO:MF | GO:0030554 | adenyl nucleotide binding | 340 | 64 | | 1.6e−11 |
| 13 | GO:MF | GO:0032559 | adenyl ribonucleotide binding | 332 | 63 | | 1.8e−11 |
| 14 | GO:MF | GO:0031489 | myosin V binding | 15 | 13 | | 2.5e−11 |
| 15 | GO:MF | GO:0008094 | DNA−dependent ATPase activity | 88 | 29 | | 1.1e−10 |
| 16 | GO:MF | GO:0140097 | catalytic activity, acting on DNA | 178 | 41 | | 9.1e−10 |
| 17 | GO:MF | GO:0003924 | GTPase activity | 250 | 49 | | 3.8e−09 |
| 18 | GO:MF | GO:0032550 | purine ribonucleoside binding | 204 | 42 | | 2.4e−08 |
| 19 | GO:MF | GO:0032549 | ribonucleoside binding | 204 | 42 | | 2.4e−08 |
| 20 | GO:MF | GO:0001883 | purine nucleoside binding | 205 | 42 | | 2.8e−08 |
| 21 | GO:MF | GO:0003724 | RNA helicase activity | 43 | 18 | | 4.1e−08 |
| 22 | GO:MF | GO:0003678 | DNA helicase activity | 59 | 21 | | 4.3e−08 |
| 23 | GO:MF | GO:0001882 | nucleoside binding | 208 | 42 | | 4.5e−08 |
| 24 | GO:MF | GO:0019001 | guanyl nucleotide binding | 217 | 43 | | 5.1e−08 |
| 25 | GO:MF | GO:0032561 | guanyl ribonucleotide binding | 216 | 42 | | 1.6e−07 |
| 26 | GO:MF | GO:0016706 | 2−oxoglutarate−dependent dioxygenase activity | 48 | 18 | | 3.6e−07 |
| 27 | GO:MF | GO:0031072 | heat shock protein binding | 95 | 24 | | 5.3e−06 |
| 28 | GO:MF | GO:0001671 | ATPase activator activity | 20 | 11 | | 5.8e−06 |
| 29 | GO:MF | GO:0050321 | tau−protein kinase activity | 20 | 11 | | 5.8e−06 |
| 30 | GO:MF | GO:0140098 | catalytic activity, acting on RNA | 330 | 50 | | 2.3e−05 |
| 31 | GO:MF | GO:0004683 | calmodulin−dependent protein kinase activity | 19 | 10 | | 4.8e−05 |
| 32 | GO:MF | GO:0000287 | magnesium ion binding | 146 | 29 | | 5.4e−05 |
| 33 | GO:MF | GO:0060590 | ATPase regulator activity | 34 | 13 | | 6.5e−05 |
| 34 | GO:MF | GO:0051787 | misfolded protein binding | 24 | 11 | | 6.7e−05 |
| 35 | GO:MF | GO:0051082 | unfolded protein binding | 93 | 22 | | 7.7e−05 |
| 36 | GO:MF | GO:0051213 | dioxygenase activity | 67 | 18 | | 1.3e−04 |
| 37 | GO:MF | GO:0005525 | GTP binding | 191 | 33 | | 2.3e−04 |
| 38 | GO:MF | GO:0015036 | disulfide oxidoreductase activity | 32 | 12 | | 2.6e−04 |
| 39 | GO:MF | GO:0016667 | oxidoreductase activity, acting on a sulfur group of donors | 50 | 15 | | 2.8e−04 |
| 40 | GO:MF | GO:0051087 | chaperone binding | 86 | 20 | | 3.8e−04 |
| 41 | GO:MF | GO:0030332 | cyclin binding | 29 | 11 | | 6.8e−04 |
| 42 | GO:MF | GO:0048407 | platelet−derived growth factor binding | 11 | 7 | | 8.1e−04 |
| 43 | GO:MF | GO:0000405 | bubble DNA binding | 8 | 6 | | 1.2e−03 |
| 44 | GO:MF | GO:0044183 | protein folding chaperone | 20 | 9 | | 1.2e−03 |
| 45 | GO:MF | GO:0001228 | DNA−binding transcription activator activity, RNA polymerase II−specific | 344 | 47 | | 1.2e−03 |
| 46 | GO:MF | GO:0030544 | Hsp70 protein binding | 31 | 11 | | 1.5e−03 |
| 47 | GO:MF | GO:0001216 | DNA−binding transcription activator activity | 347 | 47 | | 1.6e−03 |
| 48 | GO:MF | GO:0044389 | ubiquitin−like protein ligase binding | 281 | 40 | | 2.5e−03 |
| 49 | GO:MF | GO:0016705 | oxidoreductase activity, acting on paired donors, with incorporation or reduction of molecular oxygen | 148 | 26 | | 2.7e−03 |
| 50 | GO:MF | GO:0031625 | ubiquitin protein ligase binding | 264 | 38 | | 3.3e−03 |
| 51 | GO:MF | GO:0003899 | DNA−directed 5'−3' RNA polymerase activity | 28 | 10 | | 3.9e−03 |
| 52 | GO:MF | GO:0009378 | four−way junction helicase activity | 6 | 5 | | 4.0e−03 |
| 53 | GO:MF | GO:0008353 | RNA polymerase II CTD heptapeptide repeat kinase activity | 18 | 8 | | 4.9e−03 |
| 54 | GO:MF | GO:0061630 | ubiquitin protein ligase activity | 257 | 36 | | 1.1e−02 |
| 55 | GO:MF | GO:0017022 | myosin binding | 58 | 14 | | 1.1e−02 |
| 56 | GO:MF | GO:0097747 | RNA polymerase activity | 31 | 10 | | 1.1e−02 |
| 57 | GO:MF | GO:0034062 | 5'−3' RNA polymerase activity | 31 | 10 | | 1.1e−02 |
| 58 | GO:MF | GO:0043138 | 3'−5' DNA helicase activity | 16 | 7 | | 2.1e−02 |
| 59 | GO:MF | GO:0008301 | DNA binding, bending | 16 | 7 | | 2.1e−02 |
| 60 | GO:MF | GO:0048156 | tau protein binding | 40 | 11 | | 2.3e−02 |
| 61 | GO:MF | GO:0061659 | ubiquitin−like protein ligase activity | 267 | 36 | | 2.4e−02 |
| 62 | GO:MF | GO:0008474 | palmitoyl−(protein) hydrolase activity | 12 | 6 | | 3.0e−02 |
| 63 | GO:MF | GO:0015037 | peptide disulfide oxidoreductase activity | 12 | 6 | | 3.0e−02 |
| 64 | GO:MF | GO:0015035 | protein disulfide oxidoreductase activity | 12 | 6 | | 3.0e−02 |
| 65 | GO:MF | GO:0017116 | single−stranded DNA helicase activity | 17 | 7 | | 3.3e−02 |
| 66 | GO:MF | GO:0002039 | p53 binding | 57 | 13 | | 4.1e−02 |
| 67 | GO:MF | GO:0008559 | ATPase−coupled xenobiotic transmembrane transporter activity | 5 | 4 | | 5.0e−02 |

g:Profiler (biit.cs.ut.ee/gprofiler)
