## Supplemental Table 8 for "Comparative genomic and crystal structure analyses identify a collagen glucosyltransferase from *Acanthamoeba Polyphaga Mimivirus*"

| id | source | term_id | term_name | term_size | intersection_size | | p_value |
|---|---|---|---|---|---|---|---|
| 1 | REAC | REAC:R−HSA−8948216 | Collagen chain trimerization | 44 | 41 | | 1.1e−42 |
| 2 | REAC | REAC:R−HSA−8873719 | RAB geranylgeranylation | 65 | 49 | | 6.0e−42 |
| 3 | REAC | REAC:R−HSA−1650814 | Collagen biosynthesis and modifying enzymes | 67 | 48 | | 2.9e−39 |
| 4 | REAC | REAC:R−HSA−983168 | Antigen processing: Ubiquitination & Proteasome degradation | 308 | 94 | | 9.1e−36 |
| 5 | REAC | REAC:R−HSA−1474290 | Collagen formation | 89 | 48 | | 1.2e−30 |
| 6 | REAC | REAC:R−HSA−1442490 | Collagen degradation | 64 | 35 | | 3.5e−22 |
| 7 | REAC | REAC:R−HSA−2022090 | Assembly of collagen fibrils and other multimeric structures | 60 | 32 | | 1.1e−19 |
| 8 | REAC | REAC:R−HSA−8951664 | Neddylation | 233 | 61 | | 1.5e−18 |
| 9 | REAC | REAC:R−HSA−1474244 | Extracellular matrix organization | 298 | 67 | | 1.2e−16 |
| 10 | REAC | REAC:R−HSA−216083 | Integrin cell surface interactions | 84 | 32 | | 3.5e−14 |
| 11 | REAC | REAC:R−HSA−1474228 | Degradation of the extracellular matrix | 140 | 41 | | 8.2e−14 |
| 12 | REAC | REAC:R−HSA−8866652 | Synthesis of active ubiquitin: roles of E1 and E2 enzymes | 30 | 18 | | 1.1e−11 |
| 13 | REAC | REAC:R−HSA−3000178 | ECM proteoglycans | 75 | 26 | | 3.9e−10 |
| 14 | REAC | REAC:R−HSA−8876198 | RAB GEFs exchange GTP for GDP on RABs | 89 | 28 | | 7.8e−10 |
| 15 | REAC | REAC:R−HSA−8852135 | Protein ubiquitination | 78 | 26 | | 1.1e−09 |
| 16 | REAC | REAC:R−HSA−9007101 | Rab regulation of trafficking | 121 | 32 | | 3.9e−09 |
| 17 | REAC | REAC:R−HSA−186797 | Signaling by PDGF | 54 | 21 | | 4.9e−09 |
| 18 | REAC | REAC:R−HSA−419037 | NCAM1 interactions | 42 | 18 | | 2.2e−08 |
| 19 | REAC | REAC:R−HSA−3371556 | Cellular response to heat stress | 87 | 25 | | 1.1e−07 |
| 20 | REAC | REAC:R−HSA−375165 | NCAM signaling for neurite out−growth | 59 | 20 | | 2.8e−07 |
| 21 | REAC | REAC:R−HSA−3000171 | Non−integrin membrane−ECM interactions | 58 | 19 | | 1.5e−06 |
| 22 | REAC | REAC:R−HSA−3371453 | Regulation of HSF1−mediated heat shock response | 67 | 19 | | 2.1e−05 |
| 23 | REAC | REAC:R−HSA−2214320 | Anchoring fibril formation | 15 | 9 | | 4.4e−05 |
| 24 | REAC | REAC:R−HSA−8874081 | MET activates PTK2 signaling | 30 | 12 | | 1.0e−04 |
| 25 | REAC | REAC:R−HSA−73894 | DNA Repair | 328 | 47 | | 2.3e−04 |
| 26 | REAC | REAC:R−HSA−8854214 | TBC/RABGAPs | 44 | 14 | | 2.7e−04 |
| 27 | REAC | REAC:R−HSA−6782135 | Dual incision in TC−NER | 64 | 17 | | 3.0e−04 |
| 28 | REAC | REAC:R−HSA−5696398 | Nucleotide Excision Repair | 109 | 23 | | 3.1e−04 |
| 29 | REAC | REAC:R−HSA−203927 | MicroRNA (miRNA) biogenesis | 23 | 10 | | 4.3e−04 |
| 30 | REAC | REAC:R−HSA−3371571 | HSF1−dependent transactivation | 24 | 10 | | 6.9e−04 |
| 31 | REAC | REAC:R−HSA−3000157 | Laminin interactions | 30 | 11 | | 9.2e−04 |
| 32 | REAC | REAC:R−HSA−6782210 | Gap−filling DNA repair synthesis and ligation in TC−NER | 63 | 16 | | 1.2e−03 |
| 33 | REAC | REAC:R−HSA−499943 | Interconversion of nucleotide di− and triphosphates | 27 | 10 | | 2.5e−03 |
| 34 | REAC | REAC:R−HSA−8875878 | MET promotes cell motility | 40 | 12 | | 3.5e−03 |
| 35 | REAC | REAC:R−HSA−168164 | Toll Like Receptor 3 (TLR3) Cascade | 92 | 19 | | 4.1e−03 |
| 36 | REAC | REAC:R−HSA−6804756 | Regulation of TP53 Activity through Phosphorylation | 92 | 19 | | 4.1e−03 |
| 37 | REAC | REAC:R−HSA−2243919 | Crosslinking of collagen fibrils | 18 | 8 | | 4.6e−03 |
| 38 | REAC | REAC:R−HSA−6781827 | Transcription−Coupled Nucleotide Excision Repair (TC−NER) | 77 | 17 | | 4.8e−03 |
| 39 | REAC | REAC:R−HSA−450341 | Activation of the AP−1 family of transcription factors | 10 | 6 | | 7.3e−03 |
| 40 | REAC | REAC:R−HSA−3000480 | Scavenging by Class A Receptors | 19 | 8 | | 7.4e−03 |
| 41 | REAC | REAC:R−HSA−166166 | MyD88−independent TLR4 cascade | 96 | 19 | | 7.8e−03 |
| 42 | REAC | REAC:R−HSA−937061 | TRIF(TICAM1)−mediated TLR4 signaling | 96 | 19 | | 7.8e−03 |
| 43 | REAC | REAC:R−HSA−5633007 | Regulation of TP53 Activity | 160 | 26 | | 9.8e−03 |
| 44 | REAC | REAC:R−HSA−8866654 | E3 ubiquitin ligases ubiquitinate target proteins | 58 | 14 | | 9.9e−03 |
| 45 | REAC | REAC:R−HSA−187687 | Signalling to ERKs | 31 | 10 | | 1.0e−02 |
| 46 | REAC | REAC:R−HSA−171007 | p38MAPK events | 11 | 6 | | 1.5e−02 |
| 47 | REAC | REAC:R−HSA−5621481 | C−type lectin receptors (CLRs) | 136 | 23 | | 1.5e−02 |
| 48 | REAC | REAC:R−HSA−6806834 | Signaling by MET | 76 | 16 | | 1.6e−02 |
| 49 | REAC | REAC:R−HSA−5651801 | PCNA−Dependent Long Patch Base Excision Repair | 21 | 8 | | 1.8e−02 |
| 50 | REAC | REAC:R−HSA−382556 | ABC−family proteins mediated transport | 103 | 19 | | 2.2e−02 |
| 51 | REAC | REAC:R−HSA−450294 | MAP kinase activation | 63 | 14 | | 2.7e−02 |
| 52 | REAC | REAC:R−HSA−174417 | Telomere C−strand (Lagging Strand) Synthesis | 28 | 9 | | 2.7e−02 |
| 53 | REAC | REAC:R−HSA−73933 | Resolution of Abasic Sites (AP sites) | 36 | 10 | | 4.3e−02 |
| 54 | REAC | REAC:R−HSA−1296072 | Voltage gated Potassium channels | 43 | 11 | | 4.4e−02 |
| 55 | REAC | REAC:R−HSA−6805567 | Keratinization | 215 | 30 | | 4.6e−02 |

g:Profiler (biit.cs.ut.ee/gprofiler)
