## Supplemental Table 12 for "Comparative genomic and crystal structure analyses identify a collagen glucosyltransferase from *Acanthamoeba Polyphaga Mimivirus*"

**Table 1. Data collection and refinement statistics.**

| | R699 |
|---|---|
| Wavelength | |
| Resolution range | 52.67 - 2.483 (2.572 - 2.483) |
| Space group | P 21 21 21 |
| Unit cell | 84.4419 91.0219 134.762 90 90 90 |
| Total reflections | 240323 (23102) |
| Unique reflections | 37075 (3509) |
| Multiplicity | 6.5 (6.6) |
| Completeness (%) | 98.94 (95.02) |
| Mean I/sigma(I) | 26.65 (3.74) |
| Wilson B-factor | 37.51 |
| R-merge | 0.1778 (0.5341) |
| R-meas | 0.194 (0.5804) |
| R-pim | 0.07637 (0.2247) |
| CC1/2 | 0.948 (0.654) |
| CC* | 0.986 (0.889) |
| Reflections used in refinement | 36986 (3494) |
| Reflections used for R-free | 1867 (180) |
| R-work | 0.2836 (0.4819) |
| R-free | 0.3214 (0.4184) |
| CC(work) | 0.848 (0.127) |
| CC(free) | 0.791 (-0.009) |
| Number of non-hydrogen atoms | 7472 |
| macromolecules | 7326 |
| ligands | 88 |
| solvent | 58 |

| | |
|---|---|
| **Protein residues** | 891 |
| **RMS(bonds)** | 0.007 |
| **RMS(angles)** | 1.39 |
| **Ramachandran favored (%)** | 95.04 |
| **Ramachandran allowed (%)** | 4.74 |
| **Ramachandran outliers (%)** | 0.23 |
| **Rotamer outliers (%)** | 7.66 |
| **Clashscore** | 12.82 |
| **Average B-factor** | 52.15 |
| **macromolecules** | 52.36 |
| **ligands** | 50.07 |
| **solvent** | 28.91 |

Statistics for the highest-resolution shell are shown in parentheses.
